# VarStack: a Web Tool for Data Retrieval to Interpret Somatic Variants in Cancer

**DOI:** 10.1101/2020.03.10.985952

**Authors:** Morgan Howard, Bruce Kane, Mary Lepry, Paul Stey, Ashok Ragavendran, Ece D. Gamsiz Uzun

## Abstract

**Background and objective:** Advances in tumor genome sequencing created an urgent need for bioinformatics tools to support the interpretation of the clinical significance of the variants detected. VarStack is a web tool which is a base to retrieve somatic variant data in cancer from existing databases.

**Methods:** VarStack incorporates data from several publicly available databases and presents them with an easy-to-navigate user-interface. It currently supports data from the Catalogue of Somatic Mutations in Cancer (COSMIC), gnomAD, cBioPortal, ClinVar, OncoKB and UCSC Genome browser. It retrieves the data from these databases and returns back to the user in a fraction of the time it would take to manually navigate each site independently.

**Results:** Users submit a variant with gene symbol, peptide change, and coding sequence change. They may select a variety of tumor specific studies in cBioportal to search through in addition to their original query. The results from the databases are presented in tabs. Users can export the results as a CSV file. VarStack also has the batch search feature in which user submits a list of variants and download a CSV file with the data from the databases. With the batch search and data download options users can easily incorporate VarStack into their workflow or tools. VarStack saves time by providing variant data to the user from multiple databases in an easy-to-export and interpretable format.

**Availability:** VarStack is freely available under https://varstack.brown.edu.

## Introduction

Advances in sequencing technologies and bioinformatics have improved precision medicine approaches by identifying tumor-specific variants and targeting those with treatments specific to the patients. As a result, tumor variant data is large and growing. Interpretation of the variants detected is a crucial step for clinical decision-making. However, it could be time consuming as there are multiple databases available providing information on the variants. Most cancer institutions or hospitals have in-house tools to retrieve variant data. Those tools needed to be updated with the recent versions of the databases. An easy-to-interpret and user-friendly tool with up-to-date information would be beneficial for rapid clinical decision-making.

Catalogue of Somatic Mutations in Cancer (COSMIC)(1, 2), gnomAD (3), cBioPortal (4, 5), ClinVar (6), OncoKB (7), CiViC (8) and UCSC Genome browser (9) are among the existing databases used by physicians and scientists for variant interpretation. We developed VarStack 1.0 in order to provide data from the available databases on a fast and easy-to-interpret platform.

## VarStack

VarStack is a webtool which retrieves up-to-date variant information from COSMIC, gnomAD, cBioPortal, ClinVar, OncoKB and UCSC Genome browser and displays the output in separate tabs. It takes the gene name, amino acid and nucleotide change of a variant as an input. The COSMIC tab provides COSMIC ID, FATHMM prediction, coordinates in hg38 and Ensembl ID (1, 2). ClinVar tab includes the type of variant, clinical significance, dbSNP ID as well as a link to dbSNP for the specific variant, phenotype list, coordinates in hg19 and hg38 (6, 10). VarStack also provides minor allele frequency information from gnomAD (3). As an option, the user selects specific studies from the cBioportal list according to the tumor type (4). If this option is used, a graph with the frequency of the variant in the specific tumor type as well as a list of samples from the studies selected are provided under cBioPortal tab. The genomic location of the variant can be visualized on UCSC Genome Browser tab without navigating to the website (9). The user can review the data on the tabs or export it in CSV format. OncoKB tab in the downloadable CSV file is split into 3 different tabs, the first contains the actionable variant information, the second contains the curated gene information, and the last tab displays all of the annotated variant information.

VarStack also has a batch search feature which provides the data for a list of variants as a downloadable CSV file. The user enters the list of variants in a search box and the webtool returns a CSV file with the data in separate tabs for ClinVar, COSMIC, gnomAD and OncoKB. With the data export and batch search options, VarStack could be easily incorporated into existing tools or workflows.

Table 1 shows the duration VarStack takes compared to that to navigate through the individual databases (included in VarStack) for three variants. VarStack outputs the information in <50% of the time when the user navigates through each database separately (Table 1).

**Table 1:** Comparison of the time to navigate the cancer databases and VarStack

| Variants | Time (seconds) |  |  |  |  |  |  | VarStack |
| --- | --- | --- | --- | --- | --- | --- | --- | --- |
|  | COSMIC | ClinVar | gnomAD | cBioPortal | OncoKB | UCSC Genome Browser | Total |  |
| KRAS p.G12V c.35G>T | 25.99 | 7.82 | 19.9 | 27.29 | 7.73 | 6.42 | **95.15** | 26.4 |
| IDH1 p.R132H c.395G>A | 17.78 | 13.72 | 9.57 | 29.68 | 6.99 | 5.17 | **82.91** | 46.73 |
| EGFR p.E746_A750del<br>c.2235_2249del | 21.16 | 18.89 | 12.27 | 34.65 | 11.49 | 7.67 | **106.13** | 44.43 |

## System Description

VarStack is comprised of several different components including Dash (Python framework), Python (v.3.7.2), MySQL and R (v.3.5.3). Dash uses Flask, React.js and Plotly.js to provide a framework for designing web applications. It was chosen to manage the user interface and connect to a MySQL database that stores the up-to-date data from COSMIC (1, 2), ClinVar (6), gnomAD (3) and OncoKB (7). The genomic location of the variant can be viewed through an iframe of the UCSC browser using GRCh38 coordinates retrieved from the COSMIC data table (9). R package cgdsr, an API that accesses the Cancer Genomics Data Server (CGDS) was used to retrieve data from cBioPortal studies (4) (Figure 1).

**Figure 1:**
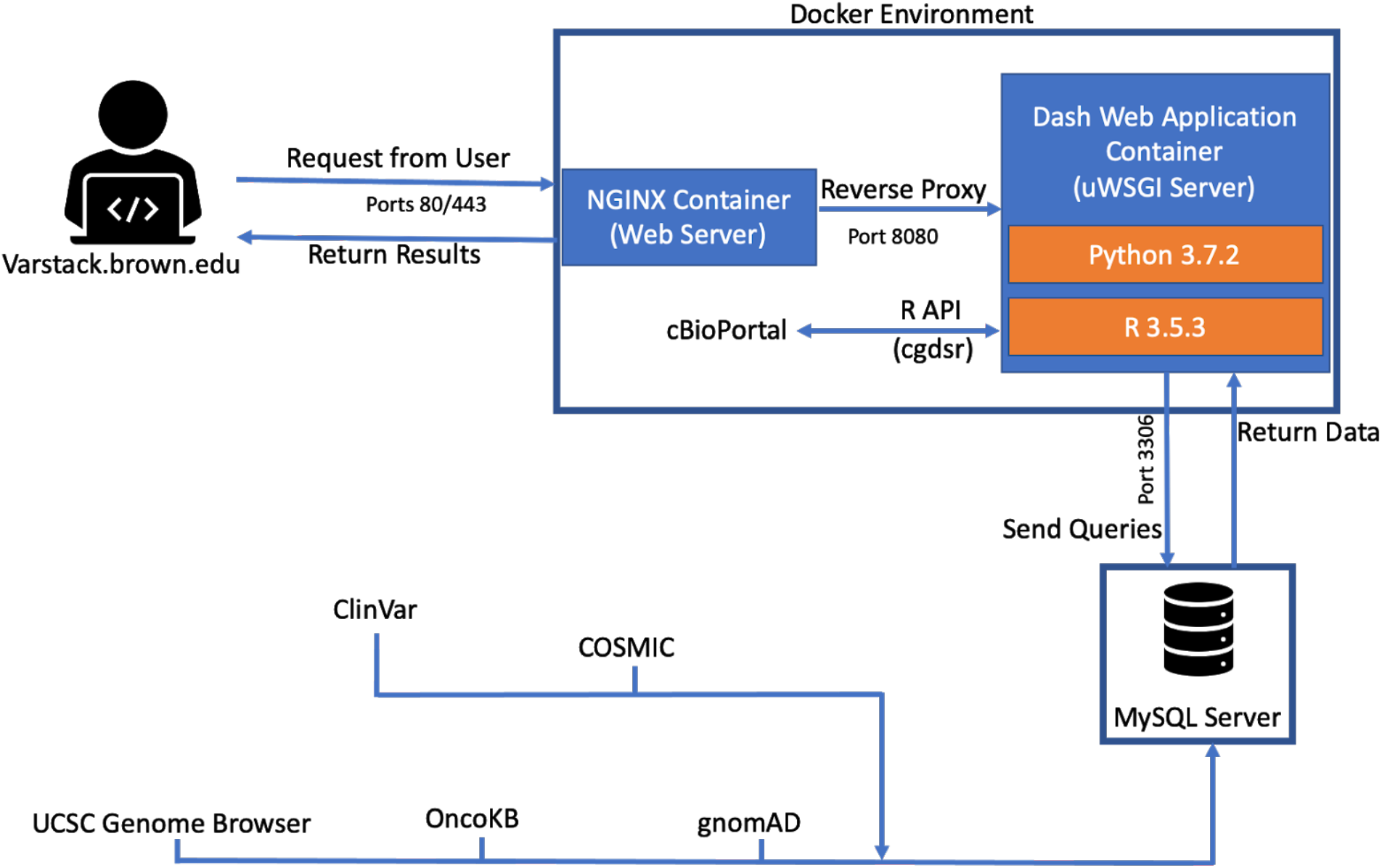
VarStack uses NGINX, R and Python to retrieve information from COSMIC, ClinVar, gnomAD, cBiportal and OncoKB.

## Discussion

The data retrieved and provided by VarStack could be obtained by manually navigating COSMIC, ClinVar, gnomAD, cBioportal, OncoKB and UCSC Genome Browser. However, this would be time-consuming in order to retrieve information for multiple variants. VarStack outputs the data in a user-friendly interface which is easy to navigate and download on one platform instead of navigating through separate websites. Data download option provides a CSV file containing the data provided in the user interface and the user does not have to save the data separately from those databases. This CSV file provides information similar to how it is displayed on VarStack in tabs with ClinVar, COSMIC, gnomAD, cBioportal and OncoKB. The batch search feature provides data from ClinVar, COSMIC, gnomAD and OncoKB for multiple variants as a downloadable CSV file. The batch search does not contain the cBioportal information as it requires specific tumor types. This feature will be implemented in the next versions. Downloadable CSV and batch search options make VarStack very compatible to be implemented for existing workflows or tools.

VarStack provides up-to-date information as it is updated according to the recent updates in the databases, but does not include expert input. The existing knowledge-base databases such as CiViC which provide expert input in addition to the information from specific databases. Although expert input is important in variant interpretation, VarStack provides up-to-date information from commonly navigated databases for somatic variant interpretation in cancer.

## Conclusions

VarStack is designed to be useful for the scientists and physicians who interpret somatic variants in multiple samples. Navigating multiple databases for several samples could be time-consuming. Also, certain databases require bioinformatics or programming knowledge. The easy-to-navigate user interface of VarStack does not require any bioinformatics or programming knowledge and saves time for the users. The tab separated user interface is a practical solution in terms of providing information from multiple databases on one base with a downloadable file and batch search features.

## Competing interests

The authors declare that they have no competing interests.

## Authors’ Contributions

M.H developed the webtool. A.R and P.S provided guidance to the infrastructure and features of the webtool. B.K and M.L helped to build the database. M.H and E.D.G.U edited the manuscript. E.D.G.U conceptualized the project and supervised the study.

## Acknowledgments

We thank Aymen Shahin as well as Brown University Computing and Information Services for their help. We also thank Dr. Alper Uzun from Brown University Alpert Medical School Department of Pediatrics for his valuable feedback.

## Notes

https://varstack.brown.edu

